# Analyses of allele age and fitness impact reveal human beneficial alleles to be older than neutral controls

**DOI:** 10.1101/2023.10.09.561569

**Authors:** Alyssa M. Pivirotto, Alexander Platt, Ravi Patel, Sudhir Kumar, Jody Hey

## Abstract

A classic population genetic prediction is that alleles experiencing directional selection should swiftly traverse allele frequency space, leaving detectable reductions in genetic variation in linked regions. However, despite this expectation, identifying clear footprints of beneficial allele passage has proven to be surprisingly challenging. We addressed the basic premise underlying this expectation by estimating the ages of large numbers of beneficial and deleterious alleles in a human population genomic data set. Deleterious alleles were found to be young, on average, given their allele frequency. However, beneficial alleles were older on average than non-coding, non-regulatory alleles of the same frequency. This finding is not consistent with directional selection and instead indicates some type of balancing selection. Among derived beneficial alleles, those fixed in the population show higher local recombination rates than those still segregating, consistent with a model in which new beneficial alleles experience an initial period of balancing selection due to linkage disequilibrium with deleterious recessive alleles. Alleles that ultimately fix following a period of balancing selection will leave a modest ‘soft’ sweep impact on the local variation, consistent with the overall paucity of species-wide ‘hard’ sweeps in human genomes.

**Impact Statement:** Analyses of allele age and evolutionary impact reveal that beneficial alleles in a human population are often older than neutral controls, suggesting a large role for balancing selection in adaptation.

## Introduction

Evolutionary adaptation depends upon the spread and fixation of beneficial alleles, however some neutral and slightly deleterious alleles also drift to high frequencies and become fixed, and so investigators have long sought ways to distinguish the fixation processes of adaptive alleles from those that are non-adaptive. Most methods are based on the classic population genetic prediction that beneficial alleles should move quickly through the range of allele frequencies (3, 4) and leave a significant footprint on levels and patterns of linked variation (5). However, despite evidence that the fixation of beneficial alleles is common (6–8), investigators have found few instances where individual fixation events have left a clear footprint (9–11). In the human context, this has been particularly puzzling given that other methods suggest that there have been thousands of adaptive amino-acid substitutions in the human lineage since the common ancestor with chimpanzees (6, 12–15).

Consequently, much research in recent years has been devoted to understanding the fixation process of beneficial alleles and the kinds of impacts that may be left in contexts of multiple mutations (16, 17), changing selection coefficients (18), selection at linked sites (19), and population structure (20–22).

To better understand the allele frequency trajectories of beneficial alleles, we undertook a new kind of analysis that combines two unrelated advances of recent years, one that can identify a large number of segregating beneficial and deleterious alleles, and another that estimates allele age. Our initial goal was to test the fundamental population genetic prediction that alleles under directional selection should be younger, on average, than neutral alleles of the same frequency. This expectation was clearly affirmed for candidate deleterious alleles; however, the analysis revealed a striking pattern in which candidate beneficial alleles are older on average than neutral alleles.

For nonsynonymous single nucleotide polymorphisms (SNPs) in a whole-genome sequencing study of over 3600 individuals from the United Kingdom (23), we identified candidate alleles under selection using the evolutionary probability (EP) of amino acids residing at each position in 17,209 autosomal genes calculated from a multi-species protein sequence alignment (24). EP estimates are based on alignments of a large number of vertebrate genomes and do not depend on the alleles currently segregating in a population or their frequency. The use of EP estimates for identifying alleles under selection is well supported by simulation (25), and they are increasingly used to identify nonsynonymous changes that are candidates for adaptive changes (26–30). As shown in Figure 1A, EP values correlate with allele frequency, with common alleles tending to have higher EP values as expected if high EP alleles are favored by selection more than are low EP alleles.

**Figure 1.**
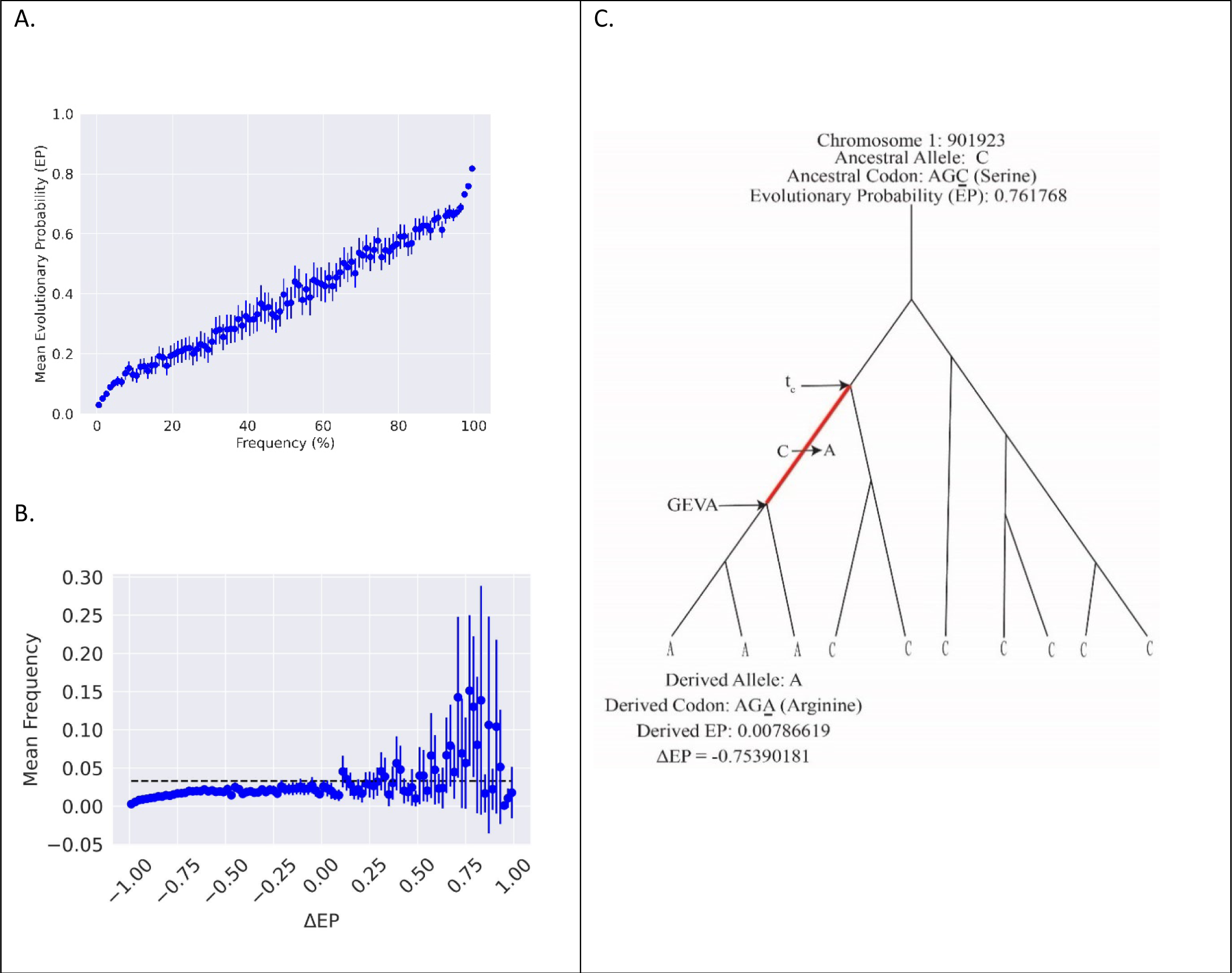
A. EP for non-synonymous SNPs binned by allele frequency. Both alleles of each SNP are included. Each bin includes a 95% confidence interval on the mean. Sites with higher EP are found at a higher frequency on average while sites with lower EP are found at lower frequencies. B. Mean derived-allele frequency binned by ΔEP values. Each bin includes a 95% confidence interval on the mean. Dotted line represents the average frequency of a neutral (non-coding, non-regulatory) site. Higher positive ΔEP bins have a higher frequency on average as expected if these sites are beneficial. C. EP calculation and age estimation targets for GEVA and *t*_*c*_ for a hypothetical site with three copies of the derived allele in a sample of 10 genomes.

We rooted non-synonymous variants using the inferred ancestral sequence from Ensembl (1) and a maximum likelihood estimator. We defined ΔEP as the derived allele EP minus the ancestral allele EP. The large majority of derived alleles are at low frequency, as expected from basic theory (31), and we observed that mean derived allele frequency increases for sites with higher positive ΔEP (Figure 1B), as expected if they are favored by natural selection (32, 33).

To consider the ages of alleles predicted to be under directional selection, we used a large control set of non-coding, non-regulatory SNPs. These will necessarily have experienced similar mutational and recombinational processes, as well as the same demographic history, that non-synonymous SNPs have experienced, and they offer the ideal landscape upon which to inquire of the impact of selection on allele age.

## Results & Discussion

### Summary of segregating and fixed derived nonsynonymous alleles

With many rooted segregating and fixed SNPs, we can examine some basic expectations of positive and negative directional selection on non-synonymous mutations (Table 1, Supplemental Table 4). First, if adaptation operates primarily at the margins of optimality, then more non-synonymous variants will be harmful than beneficial, and the magnitude of effect for deleterious mutations should be greater on average than for beneficial mutations (34). We observe both patterns, with many more negative ΔEP alleles overall, and the mean absolute magnitude of ΔEP is much greater for negative ΔEP SNPs than for positive ΔEP SNPs (0.830 versus 0.274). Comparing fixed and segregating sites, it is expected that derived positive ΔEP alleles with a frequency of 1.0 will have larger ΔEP values than those in which both ancestral and derived alleles occur in the sample, which is confirmed (0.418 for fixed vs. 0.274 for segregating). The same prediction for negative ΔEP SNPs, with fixed alleles having a higher mean value than polymorphic alleles, was also confirmed (-0.685 vs. -0.830).

**Table 1.** ΔEP measures for fixed and polymorphic alleles.

|  | **Negative $\Delta EP$** |  | **Positive $\Delta EP$** |  |
| --- | --- | --- | --- | --- |
| **Measure** | Fixed (% or CI) | Polymorphic (% or CI) | Fixed (% or CI) | Polymorphic (% or CI) |
| **# SNPs** | 23,456 (10.4%) | 202,105 (89.6%) | 3,308 (40.4%) | 4,890 (59.6%) |
| **Mean Derived Frequency** | 1.000 | 0.023 (0.022, 0.025) | 1.000 | 0.073 (0.068, 0.080) |
| **Mean Ancestral EP** | 0.725 (0.722, 0.728) | 0.847 (0.843, 0.848) | 0.154 (0.150, 0.159) | 0.243 (0.240, 0.247) |
| **Mean Derived EP** | 0.040 (0.039, 0.041) | 0.017 (0.016, 0.018) | 0.571 (0.565, 0.579) | 0.517 (0.512, 0.522) |
| **Mean $\Delta EP$** | -0.685 (-0.689, -0.682) | -0.830 (-0.834, -0.828) | 0.418 (0.408, 0.427) | 0.274 (0.267, 0.280) |

### Deleterious mutations are younger on average while beneficial mutations are older on average than neutral mutations of the same frequency

Both positively and negatively selected alleles are expected to be younger on average than neutral alleles of the same frequency (3, 35–37). We used the Genealogical Estimation of Variant Age (GEVA) method (38) to estimate the descendent node time, or coalescent time, for genes carrying the derived allele (Figure 1C). We used RUNTC (39) to estimate *t*_*c*_, the time of the ancestral node of the edge carrying the mutation (Figure 1C). Rooted bi-allelic SNPs at non-coding, non-regulatory sites were used for a control set, identified hereafter as “neutral.” The *t*_*c*_ estimator is not a function of allele frequency, and GEVA makes only limited use of allele frequency in the setting of priors for the recombinational landscape.

Allele frequency is a strong predictor of allele age, and as expected, the mean derived-allele age rises with frequency for all three classes of SNPs (Figure 2A). For both positive and negative ΔEP SNPs, an analysis of variance (ANOVA) was conducted to test the hypothesis that selected derived alleles have the same mean age as control SNPs. In both cases the null hypothesis was strongly rejected (p = 4.17x10^-12^ for negative ΔEP SNPs and 3.04x10^-27^ for positive ΔEP SNPs). However, unlike derived negative ΔEP alleles, which were younger on average than control alleles, as predicted, the positive ΔEP SNPs are older on average. Surprisingly, across most frequency intervals, derived positive ΔEP alleles exhibit mean ages thousands of generations older than the neutral control set.

**Figure 2.**
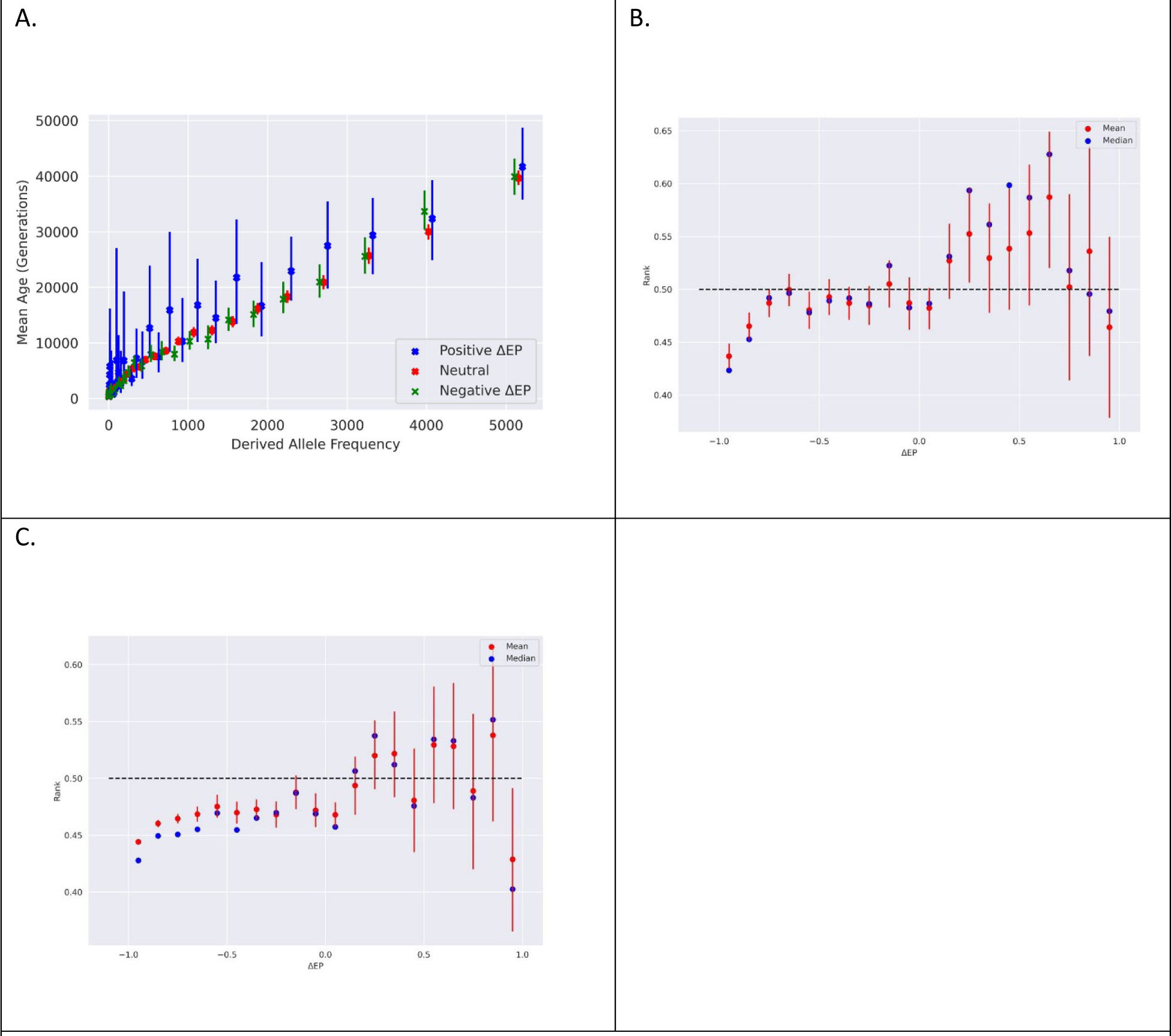
A. Allele age estimates using GEVA by allele frequency, with each frequency bin holding 75,000 neutral sites. B. Age rank (GEVA) as a function of ΔEP. Age rank for each derived allele was the rank position of the GEVA estimate in a list of all GEVA ages for neutral alleles with frequency matched derived alleles. C. Same as B, but for *t*_*c*_.

To isolate the relationship between ΔEP and allele age independently of allele frequency, we placed each allele’s age estimate into an ordered list of ages for neutral alleles of the same frequency. Non- synonymous alleles in the top half of the distribution (ranked higher than 0.5) are thus older than the median age of those neutral alleles. As shown in Figures 2B and 2C, the ranked ΔEP values show a clear trend, with negative ΔEP values falling consistently below 0.5 (i.e., with ages less than neutral alleles of the same frequency) and positive ΔEP alleles have mean age ranks consistently above 0.5.

Because the set of non-coding, non-regulatory controls necessarily experienced the same demographic context as the selected alleles, explanations of older ages for candidate beneficial alleles that depend upon interactions of selection and demography are largely ruled out, at least for models in which the beneficial alleles are indeed under directional selection. Nor can models in which these alleles are sometimes neutral and sometimes favored help explain the observation, as such alleles would still be expected to be younger on average than our control set. This pattern, in which alleles are maintained longer than alleles that are not subject to selection, is simply not consistent with positive directional selection, but rather suggests some form of balancing selection (40).

### Characterizing old, segregating, positive ΔEP alleles

Overall, a large proportion of positive ΔEP alleles are older than neutral controls. For *t*_*c*_ there were 3511 positive ΔEP alleles, 1354 of which had age ranks greater than 0.5 (38.6%). For GEVA there were 1390 positive ΔEP alleles (fewer than for *t*_*c*_ as GEVA cannot be applied to alleles that occur only once), 741 of which had age ranks greater than 0.5 (53.3%). We considered the possibility that the elevated ages of segregating positive ΔEP alleles were a kind of sampling artifact, as would occur if they represented the tail of a distribution of ages for all favored alleles, including those that became fixed (which do not appear as SNPs and for which we do not have age estimates). This explanation does not apply to alleles under strong directional selection, for which the mean and variance in sojourn times are low. On the other hand, weakly selected favored alleles will have a large mean and variance in sojourn times (41), and a large sample of such alleles would have some that, by chance, had been segregating for a long time. However, if the old segregating positive ΔEP alleles were only very weakly favored, and if they constitute the minority of alleles that were held back by the chance effects of genetic drift, then they would make up only a small fraction of all positive ΔEP alleles, including both fixed and segregating. We do not observe this in the data, with segregating alleles constituting a large fraction (0.596, Table 1) of all positive ΔEP alleles.

Balancing selection can take many forms (42), but whatever the mode of selection for these alleles, it does not appear to be the kind of long-term balancing selection that causes trans-species polymorphisms like those found in immune-related genes (43, 44). Of the positive ΔEP alleles, none of the GEVA values, and only 2.5% of the *t*_*c*_ values, are over 200,000 generations, which would correspond approximately to the human chimpanzee divergence time, assuming a 29-year generation time (45).

Most positive ΔEP sites, including those with age ranks greater than 0.5 (i.e., older than neutral alleles of the same frequency) also do not fit a conventional model of balancing selection in that the derived allele frequency is usually low (Figure 2A, Supplemental Figure 1). For *t*_*c*_ the mean frequency of positive ΔEP sites with age ranks greater than 0.5 is 0.039, while for GEVA it is 0.091.

When we seek these alleles in archaic humans, we find that relatively few positive ΔEP alleles identified in the UK10K sample (241; 4.0%) occur in a sample of 4 archaic genomes. The same analysis for negative ΔEP alleles found a smaller proportion of shared alleles (2030; 1.4%), whereas an intermediate value of noncoding sites (401,741; 3.0%) were observed among the sample of archaic genomes. For genomic regions identified as introgressed from archaics, only 13 positive ΔEP alleles (0.2% of all positive ΔEP sites) and 180 negative ΔEP alleles (0.1% of all negative ΔEP sites) were found.

We applied an alternative method for identifying balancing selection to positive ΔEP alleles that is based on the number of nearby polymorphisms that have risen to a similar frequency as the candidate allele (46). We find that the test statistic, β, is significantly higher for positive ΔEP sites compared to negative ΔEP sites (p-value = 1.588e-6), however the magnitude of these differences is small at just an average β value of 1.09 for positive ΔEP sites and 0.55 for negative ΔEP sites. Because most of the positive ΔEP sites in our study are found at low to moderate frequencies, and because the elevated ages, relative to neutral sites, are on the order of 100’s or 1000’s of generations, it is likely that there has not been sufficient time for genetic drift to bring flanking sites in to the configuration that the β statistic is designed to be sensitive to.

### Examination of modes of balancing selection: population structure and overdominance

We observed significant clumping of positive ΔEP SNPs among the genes included in the study. For every autosome, the observed variance in SNP density was significantly greater than that generated by population genetic simulation (Supplemental Table 1). Gene ontology analyses for genes rich in positive ΔEP SNPs revealed enrichment in several categories (Supplemental Table 2), most notably blood coagulation and several disease pathways.

One mechanism that could give rise to new balanced polymorphisms is if the selection regime arose because of the human population structure that favored ancestral alleles in some populations and derived alleles in other populations (as suggested in a recent analysis (47)). To examine the possibility that population structure is facilitating a large amount of balancing selection, we examined FST in the 1000 genomes data (48). Analysis of F_ST_ values in 1000 Genomes data for alleles from the UK10K samples with positive ΔEP and age ranks greater than 0.5 found no sign that these alleles show greater population structure than control alleles (Supplemental Table 3). In three comparisons, the hypothesis that F_ST_ was higher for positive ΔEP alleles that are older than expected could not be rejected by single classification Wilcoxon test in pooled African samples versus pooled European and Asian samples (p = 0.1804), pooled European versus pooled Asian samples (p = 0.5298), and Great Britain sample versus Italian sample (p = 0.7854).

Another possibility is if heterozygous positive ΔEP sites have higher fitness than homozygotes for both the ancestral and the derived alleles. To evaluate this in a way that combined the signal from all positive ΔEP alleles, we asked whether positive ΔEP alleles had higher heterozygote counts than neutral alleles of the same allele frequencies. Analyzing SNPs with at least 100 derived allele copies, we observed equal proportions of positive ΔEP sites with more heterozygotes than the neutral class, compared to fewer; and we found a mean rank for heterozygote count for positive ΔEP sites of 0.501. A one-sided *z-*test of the null hypothesis that the mean rank was equal to or less than 0.5 did not approach statistical significance (p = 0.48). This is consistent with previously published results which failed to find evidence of overdominance at deletion sites thought to be under balancing selection (49). To assess our ability to detect heterozygote advantage using counts of heterozygotes, a power analysis was conducted using simulations that mirrored the actual data set, assuming genotypes are sampled under heterozygote advantage after selection has acted. The analyses revealed that over a wide range of weak to moderate selection coefficients where the selective advantage is less than 1% (i.e., s < 0.01), that an excess of heterozygotes is unlikely to be detected given the UK10K sample size (Supplementary Table 5).

### Models that can account for a period of balancing selection

The absence of very old, derived alleles among positive ΔEP sites suggests that the balancing selection that occurs undergoes a change of character, such that balancing selection occurs for a period of time and is then followed by directional selection or no selection (i.e. genetic drift alone) leading to a loss of one or other of the alleles. If that were not the case, then we would not expect the absence of very old alleles in this data set. To address this, we consider two models that both provide mechanisms for balancing selection and that both predict that balancing selection will be a temporary phase in the process of the fixation of beneficial alleles.

One theory to explain many positive ΔEP alleles with elevated ages includes two selection stages, including first a period of balancing selection under heterozygote advantage, after which positive directional selection carries the allele to fixation. Under this “staggered sweep” model, balancing selection occurs when a favorable allele arises on a chromosome that carries one or more recessive deleterious alleles at nearby locations, and it lasts until recombination moves the allele onto other haplotypes not having linked deleterious alleles (50). A heterozygote for this chromosomal region is initially favored because of the new allele’s dominance and the harmful allele’s recessivity, such that the net positive selection coefficient on heterozygotes is strong enough to counter the effects of genetic drift. The model is supported by the fact that individual humans, and human populations, carry very large numbers of deleterious alleles, the large majority of which are expected to be mostly recessive in their effects. Considering, for example, just loss-of-function alleles for which diploid European genomes are estimated to carry about 100 (mostly in the heterozygous state), then the odds that a new beneficial mutation arises near to, and in-phase, with a deleterious allele, may be quite high (51).

Testing the staggered sweep model is difficult because local linkage estimates, as well as *t*_*c*_ and GEVA estimates, all depend on a common estimate of the genetic map. However, we can avoid this complication, and partially test the staggered sweep model, by comparing local recombination rates near positive ΔEP alleles that are fixed to those that are segregating. If segregating alleles are under balancing selection because of linkage to deleterious alleles, and the fixed alleles include those that had escaped by recombination, we expect segregating alleles to show lower local recombination rates than fixed positive ΔEP alleles. As predicted, the recombination rates of genomic regions near fixed positive ΔEP alleles were significantly higher than for segregating alleles (Mann Whitney U test p=6.0x10^-19^, Figure 3A).

**Figure 3.**
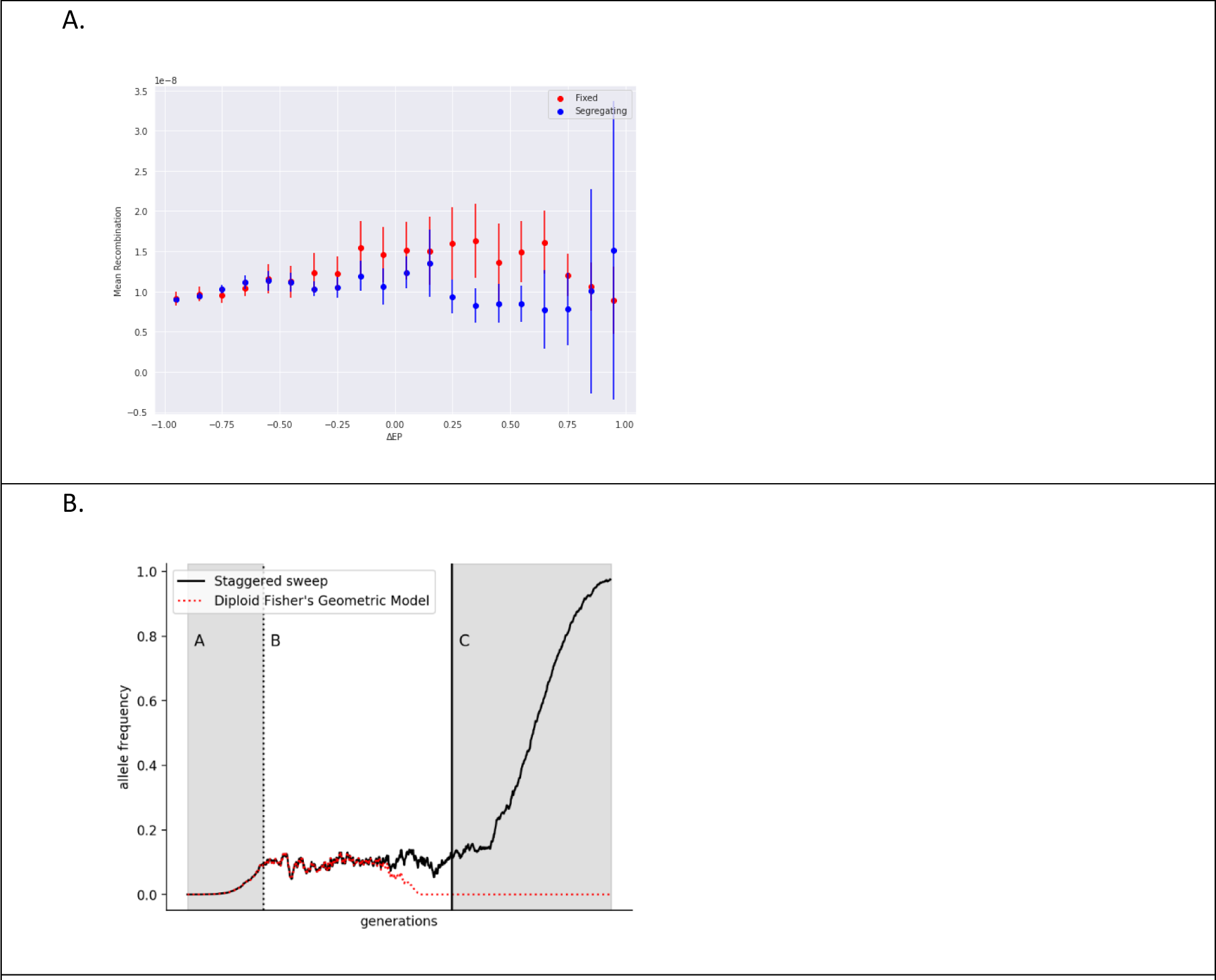
A. Mean recombination rate per base per generation as a function of ΔEP for fixed and segregating alleles. B. Figurative example of the frequency trajectory of an allele under the staggered sweep (SS) or diploid fisher’s geometric (DFG) model. Both begin with a phase of rising frequency (A) towards a period of equilibrium (B) caused by heterozygote advantage when homozygous genotypes are disfavored, either due to recessive deleterious linked variation (SS) or an overshooting of the optimal phenotype (DFG). Under DFG, variants are ultimately replaced by new mutations that are simply favored. Under SS, alleles eventually cross over onto chromosomes without linked deleterious alleles, and then rise to fixation (C).

Another explanation that also invokes heterozygote advantage is a diploid version of Fisher’s geometric model (denoted hereafter as DFG) in which mutations that carry the phenotype in the direction of the optimum may be favored when heterozygous under codominance and yet disfavored in homozygotes if that phenotype is more extreme and further away from the optimum (52). Under this model, balancing selection may be a common phase during an adaptive walk toward increasing fitness, with balanced alleles ultimately being lost when new alleles under simple positive directional selection arise and become fixed. The staggered sweep model and the DFG model differ most clearly in that the former has the period of balancing selection as a phase before the fixation of the allele, whereas the latter has the balanced allele being replaced by a new allele that is simply favored by directional selection. The former model predicts that some, perhaps many, selective sweeps are actually ‘soft’ sweeps caused by the fixation of a relatively old allele. In contrast, the DFG model predicts that when a selective sweep occurs, it is a conventional sweep by a new favored allele (i.e., a ‘hard’ sweep). Both models predict partial sweeps around new alleles that arise in a balancing selection fitness scheme (Figure 3B).

### Implications for the adaptation of human populations

We find that the majority of candidate derived beneficial alleles in a human population are segregating, rather than fixed, and yet the mean ages of these alleles are older than those for derived control alleles. These relatively old SNPs do not appear to fit a classical balancing selection model in that most of them are at low frequency and have age estimates almost always less than the age of the hominin branch.

The overall pattern suggests that when fixation of beneficial alleles does occur, it often follows an initial period of balancing selection.

We did not find evidence that ΔEP alleles are maintained due to commonly considered mechanisms of balancing selection such as population structure or heterozygote advantage, although the power to detect these factors was low, unless selection has been quite strong. Instead, we found support for the staggered sweep model in which beneficial alleles arise on the same haplotype as a deleterious mutation which delays them from fixing. Under a staggered sweep model, we predict that there should be differences in recombination rates between segregating and fixed alleles allowing for some alleles to escape selection from nearby deleterious which we find to be true for moderate positive ΔEP sites.

If many beneficial alleles have a lengthy period of balancing selection, before proceeding to fixation, then a significant fraction of adaptive fixations experienced by the human species (not just individual populations) will have occurred as a ‘soft’ sweep rather than a ‘hard’ sweep. This would help explain why there are few unambiguous cases of complete hard sweeps in large population genomic data sets (9, 11).

An additional implication of these findings is that the process of adaptation by human populations may be slower than basic population genetic models predict. If a significant fraction of ultimately beneficial fixed alleles undergoes a period of balancing selection, then at least at these sites, the process of adaptation is slowed and limited, not for lack of mutation, but rather by the process causing the period of balancing selection.

## Methods

### Evolutionary Probabilities, Allele Frequencies, and Data Filtering

Non-synonymous SNP sites in the UK10K dataset were identified with their corresponding transcript ID using the hg19 RefGene annotations in the UCSC table browser (53), that are based on NCBI RefSeq annotations (54), and the UK10K VCF (Variant Calling Format) files (23). For each two-allele polymorphism, the transcript IDs and site locations were used to retrieve the EP values for both the reference and alternative alleles. EP values were estimated using the method described in previous literature (24, 55) using posterior probabilities from a multispecies alignment with associated divergence times. Mutations excluded from this dataset include those with un-curated transcript IDs that have not been verified. Frequency data for the reference and alternative allele at each site was extracted directly from the VCF file. Analyses thought to be sensitive to CpG high mutability where limited to SNPs that did not occur as part of a CpG. These included analyses that utilized allele ages (Figures 2A, 2B, and 2C) as mutation rate was used as a parameter in estimating these values.

### Allele Age Estimates

To get approximate allele age estimates, we used both the time of most recent coalescence (*t*_*c*_) estimator (39) from the Hey Lab and the Genealogical Estimation of Variant Age (GEVA) estimator (38). To estimate *t*_*c*_, for each of the autosomal chromosome VCF files, first the singletons were phased by placing each singleton on the longer of the two haplotypes. Following this step, the time of coalescence was estimated (runtc.py) using the following parameters: k-range, mutation rate of 1e-8, and recombination map as HapMap Phase II genetic map for hg19 (56). To obtain GEVA (38) estimates, the VCF file for each autosomal chromosome was first parsed and converted into a binary file with corresponding marker and site files containing information per variant. GEVA values were obtained for all positive EP SNPs with more than two copies of the derived allele. GEVA estimates were obtained using the default parameters of effective population size of 10000, mutation rate of 1e^-8^, and the provided Hidden Markov Model (HMM) probability files. The output estimated age files were then filtered using the provided program in R (https://github.com/pkalbers/geva).

GEVA estimates were obtained for all positive ΔEP sites in the sampled genes (2729 in total). Because of time constraints large random samples of sites were used for non-coding, non-regulatory sites (71628 in total) and negative ΔEP sites (19053 in total). To generate figures with binned ΔEP values, the number of sampled noncoding, non-regulatory sites range from 800 to 2500 sites with estimated ages. For the negative ΔEP bins have approximately 1000 to 4000 sites with estimated ages, while the positive ΔEP bins have 60 to 600 with estimated ages.

### Rooting

Two methods of rooting were used, a parsimony-based approach using Ensembl (57) and a maximum likelihood approach using RAxML (58). For the parsimony-based rooting method, estimates of the hg19 ancestral states were retrieved from Ensembl (1) and included for each position in the dataset. For all analyses of allele age, SNPs were limited to those where the ancestral allele state matched the reference allele. For maximum likelihood rooting a primate alignment was extracted for each RefSeq annotated gene from an Ensembl alignment whole genome alignment (http://ftp.ensembl.org/pub/release-104/maf/ensembl-compara/multiple_alignments/12_primates.epo/12_primates.epo.10_1.maf.gz) (57).

The phylogeny for each gene was estimated using RAxML-NG using the model GTR+Γ. At positions in each gene where there was a non-synonymous mutation in the UK10K dataset, the human sequence base in the alignment was replaced with a missing value, N. Using this newly constructed primate alignment with the modified human sequence to reflect UK10K mutations, RAxML-NG was run again to estimate the base pair values at the base of the edge of the human sequence. The output generated posterior probability estimates for each of the four nucleotides at each non-synonymous SNP site. Using the posterior probabilities, the most likely ancestral state was predicted as the base pair with the highest probability. Downstream analyses were filtered by those sites where a single base pair has a probability above 0.9 indicating a higher certainty for the ancestral state.

### Calculating ΔEP

Values of ΔEP were calculated by finding the difference between the derived EP value and the ancestral EP value for a position given an estimated ancestral state for that position. The ΔEP metric indicates the difference from neutrality at a given site between the ancestral and derived allele. Sites where the amino acid mutated from an unlikely state evolutionarily to a more likely state yielded a positive ΔEP value, and in the reverse, sites where the amino acid mutated from a more likely state to less likely state yielded a negative ΔEP value.

### Noncoding variants as neutral controls

To account for allele frequency in our analyses of age across the spectrum of ΔEP values, a method to report age in relation to similar frequency control variants was needed. To assess whether an allele was young or old, each allele was compared to a large control set of alleles of the same frequency. For this purpose, we used the ages of noncoding, non-regulatory alleles, treating them as a neutral control set. Candidate SNPs for the control set were first identified from intergenic regions using annotations from SNPeff Human Genome build GRCH37 Ensembl release 75 (59, 60). This set was then filtered to remove those in regulatory regions, identified as falling into at least one of three data sets available from the UCSC Genome Browser: Candidate cis-Regulatory Elements by ENCODE (61); RefSeq Functional Elements (62); and curated regulatory annotations in the ORegAnno database (63).

Noncoding alleles in non-regulatory regions were assembled into bins of a similar frequency. Of the variants that have identified ancestral states matching the reference allele, noncoding, non-regulatory variants were split into bins of approximately 75,000 variants per frequency bin. At the lower end of frequency bins (k = 1, 2, 3, 4, 5, 6, 7, 8), same k value variants were kept together even if this resulted in bins larger than a size of 75,000 variants. In higher frequency bins, several k values were binned together to yield bins of an approximate size of 75,000 noncoding, nonregulatory variants.

### ANOVA

To test the hypotheses that neutral derived allele ages have the same mean as either beneficial or deleterious alleles we used two-way ANOVA, with selected vs control as one effect, and allele frequency bin as a second effect. We first applied the Box-Cox transformation (64) to GEVA estimates of allele age for each treatment and allele frequency group.

### Rank Analysis

To account for differences in allele ages between different frequency bins and to compare variants across the genome, we implemented a ranking system to assign each variant a rank within their own null frequency distribution. Initially, null distributions of noncoding, nonregulatory variant ages were constructed as described above. For each non-synonymous variant remaining in the filtered dataset, the corresponding frequency bin was identified based on the k value of the derived allele at that site. Within the null distribution of ages that correlated to the frequency bin for the focal non-synonymous mutation, the position of the focal mutation’s age within the null distribution was found. Based on that position, the rank within the null distribution was calculated as the position divided by the length of the null distribution (approximately 75,000 variants). This yielded a corresponding rank for each non-synonymous variant based on its own specific null distribution of ages from similar frequency variants.

### Recombination Analysis

To identify the changes in recombination across the genome, we found associated recombination rate values for every segregating and fixed non-synonymous site in the UK10K dataset. With all segregating and fixed non-synonymous sites identified using the rooting method described above, the recombination rate at that location was extracted from the genetic map file for the specific demographic in the dataset. In this case, a UK population specific recombination map (65) was used. With each site’s associated recombination rate, comparisons were made between both fixed and segregating sites across the spectrum of ΔEP values.

### F_ST_ Analysis

We examined the relationship between F_ST_ and ΔEP. In 1000 Genomes data (48), F_ST_ was calculated (66) for SNPs also found in the UK10K sample for three comparisons: pooled African samples versus pooled European and Asian samples, pooled European versus pooled Asian samples, and Great Britain sample versus Italian sample. Only SNPs with at least 10 copies of the derived allele in the pooled contrast populations were considered. Supplemental Table 3 shows mean F_ST_ as a function of ΔEP for each contrast.

To test whether F_ST_ was higher for older positive ΔEP SNPs than for control SNPs of the same allele frequencies, the F_ST_ for each positive ΔEP SNP with age rank greater than 0.5 was placed in the ranking of F_ST_ for all control SNPs of the same derived allele frequency. A single classification Wilcoxon test was conducted on each contrast to test whether there was an excess of positive ΔEP SNPs with F_ST_ ranking above 0.5.

### Heterozygosity Analysis

A test was conducted for the hypothesis that positive ΔEP SNPs have higher heterozygosity than control SNPs of the same allele frequency. For each positive ΔEP SNP, the rank position of the observed count of the number of heterozygotes was determined by placing the observed count into a sorted list of heterozygote counts for controls SNPs with the same derived allele frequency. In case of ties, the rank position was a random value of all possible ranks with the same heterozygote count. To test the hypothesis that positive ΔEP SNPs have a mean rank above 0.5, a one-sided *z*-test was conducted.

A power analysis was conducted by simulating data sets of the same size and distribution of allele frequencies as the actual data. For a given selection coefficient *s*, where the fitness of a heterozygote is 1+*s*, genotype frequencies were simulated using the observed allele count for each ΔEP SNPs in the data. Heterozygous counts were then placed in corresponding rankings of null distributions of heterozygous counts that were simulated for each of the observed allele frequencies of positive ΔEP SNPs. A *z*-test was conducted for each of 1000 simulated data sets for each selection coefficient. The results are shown in Supplemental Table 5.

### Dispersion Analysis

To assess whether positive ΔEP SNPs are evenly distributed among the genes for which we have EP values, we simulated tree-sequence (67) samples of 7242 UK chromosomes using STDPOPSIM (68) under an Out-of-Africa model (69) for each of the autosomes. Then for each autosome mutations were simulated for each gene on that chromosome, using each gene’s actual length and map position, at the same mean density as observed for positive ΔEP SNPS. The variance in simulated density of SNPs was recorded for each of 200 simulations for each autosome.

### Gene Ontology Analysis

To test whether positive ΔEP SNPs appeared more often in specific molecular, biological, and cellular classes (GO database released 2022-07-01, DOI: 10.5281/zenodo.6799722), PANTHER pathways (70) and protein classes (version 17.0, released 2022-02-22), and Reactome Pathways (Reactome database version 77, released 2021-10-01), a PANTHER Overrepresentation Test (Release 20221013) was used (71, 72). The analyzed set of genes were identified by counting the number of positive ΔEP SNPs per gene. The number of positive ΔEP SNPs was normalized by gene length, and all genes with more than one positive ΔEP were retained. A final subset of 73 genes were used in the PANTHER GO term analysis.

For the reference list, the gene database for Homo sapiens was used. Analyses were conducted with a Fisher’s Exact test with a False Discovery Rate correction. Results are detailed in Supplemental Table 2.

### Comparison to Archaic Genomes

In order to identify whether a large proportion of our sites of interest arose prior to the speciation between modern humans and archaic humans, we examined for each site whether it was also present in any one of four archaic genomes (73–76). For each category: nonsynonymous – ΔEP, nonsynonymous + ΔEP, and neutral noncoding sites, the number of shared loci with at least one archaic genome is reported along with percent of shared sites over the number of all sites in that category.

Not only was there interest in knowing whether these sites arose prior to the speciation event, but some subset of these sites potentially could be found in both modern human genomes and archaic human genomes due to gene flow between the two species. Sites were identified as appearing in introgression regions based on S* values generated from the CEU dataset from 1000 Genomes (77) (available at https://data.mendeley.com/datasets/y7hyt83vxr/1). Sites annotated as matching in either Neanderthal or Denisovan would be included as introgression sites for our analysis.

### β (^2^) Values

For β (^2^) scores (46), the CEU standardized scores generated from 1000 Genomes data was used (available at https://zenodo.org/record/7842447). For each site in our analysis, we identified from this published dataset the Beta2 score if available. A Mann-Whitney U test was done to analyze the difference between the Beta2 values of the – ΔEP and + ΔEP distributions.

## Acknowledgments

This research was supported in part by NIH grants R01GM144468-01 to J. Hey and R35GM139540-02 to S. Kumar. A. Platt was partially funded by N.I.H. grant R35 GM134957-01 and American Diabetes Association Pathway to Stop Diabetes grant #1-19-VSN-02. This research includes calculations carried out on HPC (High Performance Computing) resources supported in part by the National Science Foundation through major research instrumentation grant number 1625061 and by the US Army Research Laboratory under contract number W911NF-16-2-0189.

## Data Availability

Tables of detailed information for nonsynonymous and noncoding variants, as well as a list of primary mRNA isoforms (in the form of RefSeq IDs) used to retrieve EP values, are available at https://bio.cst.temple.edu/~tuf29449/nolinks/Pivirotto_Balancing_Selection_info.zip.

## Author Contributions

AMP, SK, AP, and JH developed the idea for the study. RP contributed evolutionary probability values. AMP and JH conducted the study, including writing scripts and conducted the analyses. AMP and JH drafted the paper, with comments and suggestions from AP and SK.

## Supplementary Figures & Tables

Supplementary Table 1. Results of simulation-based tests of dispersion of positive ΔEP SNPs.

Supplemental Table 2. Gene ontology results

Supplemental Table 3. FST Values across ΔEP spectrum of values. Mean FST rank value for UK10K SNPs in ΔEP bins for three population contrasts. Values are for SNPs that are in the UK10K sample and occur with at least 10 derived alleles in the pooled populations of the contrast. For each ΔEP SNP the observed FST was ranked against that for control alleles of the same derived allele frequency.

Supplementary Table 4. ΔEP measures for fixed and polymorphic alleles. Based on maximum-likelihood rooting estimates of ancestral alleles (see Figure 1 for values based on Ensembl rooting). Simulated mean ΔEP was calculated for each SNP by considering all possible non- synonymous mutations and the corresponding EP value for the resulting amino acid in proportion to their mutation probabilities based on empirical estimates. 95% confidence intervals on the mean, determined by bias-corrected bootstrap, are given in parentheses.

Supplementary Table 5. Statistical power for detecting excess heterozygosity.

Supplementary Figure 1. Distributions of derived polymorphism frequency in UK10K.

Distribution of derived allele frequency for each ΔEP bin from –1 to +1 in 0.1 increments. Derived allele frequency ranges from singletons (1 copy of the derived allele) to 7241 copies (only one copy of the ancestral allele). The majority of sites are found at low frequencies across all bins

## References

1. Herrero J, Muffato M, Beal K, Fitzgerald S, Gordon L, Pignatelli M, et al. Ensembl comparative genomics resources. Database. 2016;2016.

2. Efron B, Tibshirani RJ. An introduction to the bootstrap: CRC press; 1994.

3. Maruyama T. The age of an allele in a finite population. Genet Res. 1974;23(02):137–43.

4. Kimura M, Ohta T. The average number of generations until fixation of a mutant gene in a finite population. Genetics. 1969;61:763–71.

5. Maynard Smith J, Haigh J. The hitch-hiking effect of a favourable gene. Genet Res. 1974;23:23–35.

6. Uricchio LH, Petrov DA, Enard D. Exploiting selection at linked sites to infer the rate and strength of adaptation. Nature Ecology & Evolution. 2019.

7. Galtier N. Adaptive Protein Evolution in Animals and the Effective Population Size Hypothesis. PLoS Genetics. 2016;12(1):e1005774.

8. Enard D, Messer PW, Petrov DA. Genome-wide signals of positive selection in human evolution. Genome Research. 2014;24(6):885–95.

9. Hernandez RD, Kelley JL, Elyashiv E, Melton SC, Auton A, McVean G, et al. Classic Selective Sweeps Were Rare in Recent Human Evolution. Science. 2011;331(6019):920-4.

10. Schrider DR, Kern AD. Soft sweeps are the dominant mode of adaptation in the human genome. Mol Biol Evol. 2017;34(8):1863–77.

11. Coop G, Pickrell JK, Novembre J, Kudaravalli S, Li J, Absher D, et al. The Role of Geography in Human Adaptation. PLoS Genet. 2009;5(6):e1000500.

12. 12. Consortium CSaA. Initial sequence of the chimpanzee genome and comparison with the human genome. 2005;437(7055):69–87.

13. Zhen Y, Huber CD, Davies RW, Lohmueller KE. Greater strength of selection and higher proportion of beneficial amino acid changing mutations in humans compared with mice and Drosophila melanogaster. Genome Res. 2021;31(1):110–20.

14. Huber CD, Kim BY, Marsden CD, Lohmueller KE. Determining the factors driving selective effects of new nonsynonymous mutations. Proceedings of the National Academy of Sciences. 2017;114(17):4465–70.

15. Boyko AR, Williamson SH, Indap AR, Degenhardt JD, Hernandez RD, Lohmueller KE, et al. Assessing the evolutionary impact of amino acid mutations in the human genome. PLoS Genet. 2008;4(5):e1000083.

16. Garud NR, Messer PW, Petrov DA. Detection of hard and soft selective sweeps from Drosophila melanogaster population genomic data. PLoS Genetics. 2021;17(2):e1009373.

17. Harris RB, Sackman A, Jensen JD. On the unfounded enthusiasm for soft selective sweeps II: Examining recent evidence from humans, flies, and viruses. PLoS Genetics. 2018;14(12):e1007859.

18. McCoy RC, Akey JM. Selection plays the hand it was dealt: evidence that human adaptation commonly targets standing genetic variation. Genome biology. 2017;18(1):1–4.

19. Charlesworth B, Jensen JD. Effects of Selection at Linked Sites on Patterns of Genetic Variability. Annual Review of Ecology, Evolution, and Systematics. 2021;52(1):177–97.

20. Souilmi Y, Tobler R, Johar A, Williams M, Grey ST, Schmidt J, et al. Admixture has obscured signals of historical hard sweeps in humans. Nature Ecology & Evolution. 2022:1–13.

21. Novembre J, Galvani AP, Slatkin M. The geographic spread of the CCR5 Δ32 HIV-resistance allele. PLoS Biology. 2005;3(11):e339.

22. Muktupavela RA, Petr M, Ségurel L, Korneliussen T, Novembre J, Racimo F. Modeling the spatiotemporal spread of beneficial alleles using ancient genomes. Elife. 2022;11:e73767.

23. Consortium TUK. The UK10K project identifies rare variants in health and disease. Nature. 2015;526(7571):82–90.

24. Patel R, Kumar S. On estimating evolutionary probabilities of population variants. BMC Evolutionary Biology. 2019;19(1):1–14.

25. Patel R, Scheinfeldt LB, Sanderford MD, Lanham TR, Tamura K, Platt A, et al. Adaptive landscape of protein variation in human exomes. Mol Biol Evol. 2018;35(8):2015–25.

26. Pyott SJ, van Tuinen M, Screven LA, Schrode KM, Bai J-P, Barone CM, et al. Functional, morphological, and evolutionary characterization of hearing in subterranean, eusocial African mole-rats. Curr Biol. 2020;30(22):4329–41. e4.

27. Dolatyabi S, Peighambari SM, Razmyar J. Molecular detection and analysis of beak and feather disease viruses in Iran. Frontiers in Veterinary Science. 2022;9.

28. Xu K, Kosoy R, Shameer K, Kumar S, Liu L, Readhead B, et al. Genome-wide analysis indicates association between heterozygote advantage and healthy aging in humans. BMC genetics. 2019;20(1):1–14.

29. Tian R, Pan Y, Etheridge TH, Deshmukh H, Gulick D, Gibson G, et al. Pitfalls in single clone CRISPR-Cas9 mutagenesis to fine-map regulatory intervals. Genes. 2020;11(5):504.

30. Ose NJ, Campitelli P, Patel R, Kumar S, Ozkan SB. Protein dynamics provide mechanistic insights about epistasis among common missense polymorphisms. Biophysical journal. 2023.

31. Wright S. The distribution of gene frequencies in populations. Proc Natl Acad Sci U S A. 1937;23:307–20.

32. Wright S. The Distribution of Gene Frequencies Under Irreversible Mutation. Proc Natl Acad Sci. 1938;24(7):253–9.

33. Kimura M. Genetic variability maintained in a finite population due to mutational production of neutral and nearly neutral isoalleles. Genet Res. 1968;11:247–69.

34. Fisher RA. The genetical theory of natural selection. Oxford: Clarenson Press; 1930.

35. Kimura M, Ohta T. The age of a neutral mutant persisting in a finite population. Genetics. 1973;75:199–212.

36. Slatkin M, Rannala B. Estimating Allele Age. Annual Review of Genomics and Human Genetics. 2000;1(1):225–49.

37. Kiezun A, Pulit SL, Francioli LC, van Dijk F, Swertz M, Boomsma DI, et al. Deleterious Alleles in the Human Genome Are on Average Younger Than Neutral Alleles of the Same Frequency. PLoS Genet. 2013;9(2):e1003301.

38. Albers PK, McVean G. Dating genomic variants and shared ancestry in population-scale sequencing data. PLoS Biology. 2020;18(1):e3000586.

39. Platt A, Pivirotto A, Knoblauch J, Hey J. An estimator of first coalescent time reveals selection on young variants and large heterogeneity in rare allele ages among human populations. PLoS Genetics. 2019;15(8):e1008340.

40. Dobzhansky T. Mendelism, Darwinism, and evolutionism. Proc Am Philos Soc. 1965;109(4):205–15.

41. De Sanctis B, Krukov I, de Koning A. Allele age under non-classical assumptions is clarified by an exact computational Markov chain approach. Scientific reports. 2017;7(1):1–11.

42. Dobzhansky T. Genetics of the Evolutionary Process: Columbia University Press; 1971.

43. Leffler EM, Gao Z, Pfeifer S, Ségurel L, Auton A, Venn O, et al. Multiple instances of ancient balancing selection shared between humans and chimpanzees. Science. 2013;339(6127):1578-82.

44. Bitarello BD, de Filippo C, Teixeira JC, Schmidt JM, Kleinert P, Meyer D, et al. Signatures of long-term balancing selection in human genomes. Genome biology and evolution. 2018;10(3):939–55.

45. Fenner JN. Cross-cultural estimation of the human generation interval for use in genetics-based population divergence studies. Am J Phys Anthrop. 2005;128(2):415–23.

46. Siewert KM, Voight BF. BetaScan2: Standardized statistics to detect balancing selection utilizing substitution data. Genome Biology and Evolution. 2020;12(2):3873–7.

47. Soni V, Vos M, Eyre-Walker A. A new test suggests hundreds of amino acid polymorphisms in humans are subject to balancing selection. PLoS Biology. 2022;20(6):e3001645.

48. 1000 Genomes Project Consortium. A global reference for human genetic variation. Nature. 2015;526(7571):68-74.

49. Aqil A, Speidel L, Pavlidis P, Gokcumen O. Balancing selection on genomic deletion polymorphisms in humans. Elife. 2023;12:e79111.

50. Assaf ZJ, Petrov DA, Blundell JR. Obstruction of adaptation in diploids by recessive, strongly deleterious alleles. Proceedings of the National Academy of Sciences. 2015;112(20):E2658–E66.

51. Henn BM, Botigué LR, Bustamante CD, Clark AG, Gravel S. Estimating Mutation Load in Human Genomes. Nature reviews Genetics. 2015;16(6):333–43.

52. Sellis D, Callahan BJ, Petrov DA, Messer PW. Heterozygote advantage as a natural consequence of adaptation in diploids. Proceedings of the National Academy of Sciences. 2011.

53. Karolchik D, Hinrichs AS, Furey TS, Roskin KM, Sugnet CW, Haussler D, et al. The UCSC Table Browser data retrieval tool. Nucleic Acids Res. 2004;32(suppl_1):D493-D6.

54. Pruitt KD, Tatusova T, Maglott DR. NCBI Reference Sequence (RefSeq): a curated non-redundant sequence database of genomes, transcripts and proteins. Nucleic Acids Res. 2005;33(suppl_1):D501-D4.

55. Liu L, Tamura K, Sanderford M, Gray VE, Kumar S. A Molecular Evolutionary Reference for the Human Variome. Mol Biol Evol. 2015;33(1):245–54.

56. Consortium IH. A second generation human haplotype map of over 3.1 million SNPs. Nature. 2007;449(7164):851.

57. Howe KL, Achuthan P, Allen J, Allen J, Alvarez-Jarreta J, Amode MR, et al. Ensembl 2021. Nucleic Acids Res. 2021;49(D1):D884-D91.

58. Kozlov AM, Darriba D, Flouri T, Morel B, Stamatakis A. RAxML-NG: a fast, scalable and user- friendly tool for maximum likelihood phylogenetic inference. Bioinformatics. 2019;35(21):4453–5.

59. Cunningham F, Allen JE, Allen J, Alvarez-Jarreta J, Amode M R, Armean Irina M, et al. Ensembl 2022. Nucleic Acids Res. 2021;50(D1):D988-D95.

60. Cingolani P, Platts A, Wang LL, Coon M, Nguyen T, Wang L, et al. A program for annotating and predicting the effects of single nucleotide polymorphisms, SnpEff: SNPs in the genome of Drosophila melanogaster strain w1118; iso-2; iso-3. Fly. 2012;6(2):80-92.

61. Moore JE, Purcaro MJ, Pratt HE, Epstein CB, Shoresh N, Adrian J, et al. Expanded encyclopaedias of DNA elements in the human and mouse genomes. Nature. 2020;583(7818):699-710.

62. Farrell CM, Goldfarb T, Rangwala SH, Astashyn A, Ermolaeva OD, Hem V, et al. RefSeq Functional Elements as experimentally assayed nongenic reference standards and functional interactions in human and mouse. Genome Research. 2022;32(1):175–88.

63. Lesurf R, Cotto KC, Wang G, Griffith M, Kasaian K, Jones SJ, et al. ORegAnno 3.0: a community- driven resource for curated regulatory annotation. Nucleic Acids Res. 2016;44(D1):D126–D32.

64. Box GE, Cox DR. An analysis of transformations. Journal of the Royal Statistical Society Series B: Statistical Methodology. 1964;26(2):211–43.

65. Spence JP, Song YS. Inference and analysis of population-specific fine-scale recombination maps across 26 diverse human populations. Science Advances. 2019;5(10):eaaw9206.

66. Wright S. Coefficients of inbreeding and relationship. Amer Nat. 1922;56:330–8.

67. Kelleher J, Thornton KR, Ashander J, Ralph PL. Efficient pedigree recording for fast population genetics simulation. PLOS Computational Biology. 2018;14(11):e1006581.

68. Adrion JR, Cole CB, Dukler N, Galloway JG, Gladstein AL, Gower G, et al. A community-maintained standard library of population genetic models. Elife. 2020;9.

69. Tennessen JA, Bigham AW, O’Connor TD, Fu W, Kenny EE, Gravel S, et al. Evolution and functional impact of rare coding variation from deep sequencing of human exomes. Science. 2012;337(6090):64-9.

70. Mi H, Thomas P. PANTHER pathway: an ontology-based pathway database coupled with data analysis tools. Protein networks and pathway analysis: Springer; 2009. p. 123–40.

71. Thomas PD, Ebert D, Muruganujan A, Mushayahama T, Albou LP, Mi H. PANTHER: Making genome-scale phylogenetics accessible to all. Protein Science. 2022;31(1):8–22.

72. Mi H, Muruganujan A, Huang X, Ebert D, Mills C, Guo X, et al. Protocol Update for large-scale genome and gene function analysis with the PANTHER classification system (v. 14.0). Nature protocols. 2019;14(3):703-21.

73. Mafessoni F, Grote S, de Filippo C, Slon V, Kolobova KA, Viola B, et al. A high-coverage Neandertal genome from Chagyrskaya Cave. Proceedings of the National Academy of Sciences. 2020;117(26):15132–6.

74. Prüfer K, de Filippo C, Grote S, Mafessoni F, Korlević P, Hajdinjak M, et al. A high-coverage Neandertal genome from Vindija Cave in Croatia. Science. 2017;358(6363):655-8.

75. Prufer K, Racimo F, Patterson N, Jay F, Sankararaman S, Sawyer S, et al. The complete genome sequence of a Neanderthal from the Altai Mountains. Nature. 2014;505(7481):43-9.

76. Meyer M, Kircher M, Gansauge M-T, Li H, Racimo F, Mallick S, et al. A high-coverage genome sequence from an archaic Denisovan individual. Science. 2012;338(6104):222-6.

77. Browning SR, Browning BL, Zhou Y, Tucci S, Akey JM. Analysis of human sequence data reveals two pulses of archaic Denisovan admixture. Cell. 2018;173(1):53–61. e9.

