## Supplementary Information for "Analyses of allele age and fitness impact reveal human beneficial alleles to be older than neutral controls"

For “Balancing selection is common for beneficial alleles in a human population” by Pivirotto et al

**Supplementary Table 1. Results of simulation-based tests of dispersion of positive ΔEP SNPs.**

| Chromosome | # Genes | Observed Mean SNP Density/bp | Mean simulated density | Observed Variance in Density | Mean simulated variance in density | Estimated probability of simulating variance higher than observed |
|---|---|---|---|---|---|---|
| 1 | 1849 | 0.00035 | 0.00034 | 1.21E-06 | 3.50E-07 | 0 |
| 2 | 1134 | 0.0003 | 0.0003 | 1.45E-06 | 2.72E-07 | 0 |
| 3 | 1000 | 0.00027 | 0.00027 | 8.35E-07 | 2.39E-07 | 0 |
| 4 | 695 | 0.00039 | 0.00039 | 1.70E-06 | 3.75E-07 | 0 |
| 5 | 815 | 0.00043 | 0.00043 | 2.49E-06 | 4.10E-07 | 0 |
| 6 | 950 | 0.00026 | 0.00025 | 6.85E-07 | 2.69E-07 | 0 |
| 7 | 807 | 0.00038 | 0.00038 | 1.75E-06 | 3.65E-07 | 0 |
| 8 | 603 | 0.00038 | 0.00039 | 1.93E-06 | 3.70E-07 | 0 |
| 9 | 701 | 0.0004 | 0.0004 | 1.67E-06 | 3.81E-07 | 0 |
| 10 | 667 | 0.00035 | 0.00036 | 1.67E-06 | 3.25E-07 | 0 |
| 11 | 1207 | 0.00055 | 0.00055 | 2.87E-06 | 5.76E-07 | 0 |
| 12 | 965 | 0.00025 | 0.00026 | 6.31E-07 | 2.47E-07 | 0 |
| 13 | 299 | 0.00023 | 0.00023 | 5.37E-07 | 1.95E-07 | 0 |
| 14 | 551 | 0.00023 | 0.00024 | 4.00E-07 | 2.27E-07 | 0 |
| 15 | 517 | 0.00021 | 0.00022 | 3.47E-07 | 1.83E-07 | 0 |
| 16 | 744 | 0.00057 | 0.00057 | 4.37E-06 | 5.82E-07 | 0 |
| 17 | 1052 | 0.00043 | 0.00043 | 1.80E-06 | 4.31E-07 | 0 |
| 18 | 251 | 0.00023 | 0.00023 | 3.11E-07 | 1.99E-07 | 0.055 |
| 19 | 1296 | 0.00076 | 0.00076 | 4.32E-06 | 7.38E-07 | 0 |
| 20 | 506 | 0.00035 | 0.00035 | 9.00E-07 | 3.99E-07 | 0 |
| 21 | 205 | 0.00064 | 0.00065 | 5.66E-06 | 9.64E-07 | 0 |
| 22 | 395 | 0.00049 | 0.00049 | 1.74E-06 | 4.78E-07 | 0 |

**Supplemental Table 2. Gene ontology results**

| Panther Pathways | Observed | Expected | Fold Enrichment | Raw P Value | FDR |
|---|---|---|---|---|---|
| Plasminogen activating cascade | 4 | 0.06 | 64.59 | 9.14E-07 | 7.31E-05 |
| Blood coagulation | 6 | 0.17 | 35.81 | 3.33E-08 | 5.32E-06 |

| GO Biological Process | Observed | Expected | Fold Enrichment | Raw P Value | FDR |
|---|---|---|---|---|---|
| plasminogen activation | 3 | 0.04 | 74.87 | 1.62E-05 | 1.82E-02 |
| blood coagulation, fibrin clot formation | 5 | 0.09 | 57.19 | 6.10E-08 | 3.19E-04 |
| protein activation cascade | 5 | 0.09 | 52.79 | 8.68E-08 | 3.40E-04 |
| blood coagulation | 7 | 0.63 | 11.11 | 4.08E-06 | 7.11E-03 |
| coagulation | 8 | 0.64 | 12.55 | 3.30E-07 | 1.04E-03 |
| hemostasis | 7 | 0.65 | 10.80 | 4.89E-06 | 6.97E-03 |
| positive regulation of heterotypic cell-cell adhesion | 3 | 0.05 | 54.90 | 3.60E-05 | 3.32E-02 |
| acute-phase response | 6 | 0.15 | 39.22 | 2.03E-08 | 3.18E-04 |
| acute inflammatory response | 7 | 0.3 | 23.43 | 3.39E-08 | 2.65E-04 |
| platelet aggregation | 4 | 0.16 | 25.54 | 2.54E-05 | 2.65E-02 |
| cell-cell adhesion | 10 | 1.98 | 5.06 | 3.02E-05 | 2.96E-02 |
| negative regulation of endopeptidase activity | 8 | 0.9 | 8.86 | 4.10E-06 | 6.43E-03 |
| regulation of endopeptidase activity | 10 | 1.54 | 6.51 | 3.54E-06 | 6.95E-03 |
| regulation of peptidase activity | 11 | 1.65 | 6.68 | 8.55E-07 | 1.91E-03 |
| regulation of proteolysis | 14 | 2.71 | 5.17 | 4.91E-07 | 1.28E-03 |
| negative regulation of peptidase activity | 8 | 0.94 | 8.55 | 5.29E-06 | 6.91E-03 |
| negative regulation of proteolysis | 9 | 1.26 | 7.16 | 5.33E-06 | 6.43E-03 |

| GO Molecular Function | Observed | Expected | Fold Enrichment | Raw P Value | FDR |
|---|---|---|---|---|---|
| serine-type endopeptidase inhibitor activity | 6 | 0.37 | 16.31 | 2.52E-06 | 2.51E-03 |
| endopeptidase inhibitor activity | 8 | 0.67 | 12.00 | 4.57E-07 | 1.14E-03 |
| endopeptidase regulator activity | 8 | 0.72 | 11.15 | 7.81E-07 | 9.72E-04 |
| peptidase regulator activity | 9 | 0.86 | 10.51 | 2.46E-07 | 1.22E-03 |
| peptidase inhibitor activity | 8 | 0.69 | 11.56 | 6.01E-07 | 9.96E-04 |
| protease binding | 6 | 0.51 | 11.68 | 1.57E-05 | 1.30E-02 |

| GO Cellular Component | Observed | Expected | Fold Enrichment | Raw P Value | FDR |
|---|---|---|---|---|---|
| fibrinogen complex | 4 | 0.03 | > 100 | 7.75E-08 | 1.98E-05 |
| extracellular space | 26 | 12.49 | 2.08 | 1.41E-04 | 1.92E-02 |
| extracellular region | 30 | 16.01 | 1.87 | 3.16E-04 | 3.23E-02 |
| endocytic vesicle lumen | 3 | 0.08 | 37.43 | 9.97E-05 | 1.57E-02 |
| platelet alpha granule lumen | 7 | 0.24 | 28.68 | 9.23E-09 | 3.77E-06 |
| platelet alpha granule | 7 | 0.33 | 21.12 | 6.62E-08 | 1.93E-05 |
| secretory granule | 13 | 3.2 | 4.06 | 1.75E-05 | 2.98E-03 |
| secretory vesicle | 13 | 3.83 | 3.40 | 1.07E-04 | 1.57E-02 |
| secretory granule lumen | 12 | 1.17 | 10.29 | 2.56E-09 | 1.74E-06 |
| cytoplasmic vesicle lumen | 12 | 1.18 | 10.20 | 2.83E-09 | 1.45E-06 |
| vesicle lumen | 13 | 1.18 | 10.98 | 2.43E-10 | 2.48E-07 |
| blood microparticle | 13 | 0.52 | 24.78 | 1.43E-14 | 2.93E-11 |
| endoplasmic reticulum lumen | 11 | 1.14 | 9.62 | 2.46E-08 | 8.38E-06 |
| collagen-containing extracellular matrix | 11 | 1.58 | 6.96 | 5.79E-07 | 1.31E-04 |
| extracellular matrix | 11 | 2.09 | 5.25 | 8.22E-06 | 1.68E-03 |
| external encapsulating structure | 11 | 2.1 | 5.24 | 8.35E-06 | 1.55E-03 |
| extracellular exosome | 19 | 7.65 | 2.48 | 1.52E-04 | 1.94E-02 |
| extracellular vesicle | 19 | 7.73 | 2.46 | 1.75E-04 | 2.10E-02 |
| extracellular membrane-bounded organelle | 19 | 7.74 | 2.46 | 1.76E-04 | 1.89E-02 |
| extracellular organelle | 19 | 7.74 | 2.46 | 1.76E-04 | 2.00E-02 |

| Panther Protein Class | Observed | Expected | Fold Enrichment | Raw P Value | FDR |
|---|---|---|---|---|---|
| protease inhibitor | 6 | 0.48 | 12.57 | 1.05E-05 | 2.06E-03 |

| Reactome Pathways | Observed | Expected | Fold Enrichment | Raw P Value | FDR |
|---|---|---|---|---|---|
| LDL remodeling | 2 | 0.02 | 91.51 | 3.59E-04 | 2.98E-02 |
| Plasma lipoprotein remodeling | 3 | 0.12 | 25.74 | 2.76E-04 | 2.38E-02 |
| Plasma lipoprotein assembly, remodeling, and clearance | 4 | 0.24 | 16.39 | 1.29E-04 | 1.29E-02 |
| GRB2:SOS provides linkage to MAPK signaling for Integrins | 4 | 0.05 | 78.43 | 4.71E-07 | 2.35E-04 |

| | | | | | |
|---|---|---|---|---|---|
| Integrin signaling | 4 | 0.09 | 42.23 | 4.08E-06 | 1.02E-03 |
| Platelet Aggregation (Plug Formation) | 4 | 0.14 | 28.90 | 1.61E-05 | 3.35E-03 |
| Platelet activation, signaling and aggregation | 9 | 0.95 | 9.50 | 5.57E-07 | 1.98E-04 |
| p130Cas linkage to MAPK signaling for integrins | 4 | 0.05 | 73.21 | 5.95E-07 | 1.85E-04 |
| Regulation of TLR by endogenous ligand | 4 | 0.08 | 52.29 | 1.91E-06 | 5.29E-04 |
| MyD88 deficiency (TLR2/4) | 3 | 0.06 | 48.44 | 5.01E-05 | 5.94E-03 |
| Diseases associated with the TLR signaling cascade | 3 | 0.11 | 26.57 | 2.53E-04 | 2.34E-02 |
| Diseases of Immune System | 3 | 0.11 | 26.57 | 2.53E-04 | 2.43E-02 |
| IRAK4 deficiency (TLR2/4) | 3 | 0.07 | 45.75 | 5.83E-05 | 6.60E-03 |
| Common Pathway of Fibrin Clot Formation | 3 | 0.08 | 37.43 | 9.97E-05 | 1.08E-02 |
| Formation of Fibrin Clot (Clotting Cascade) | 5 | 0.14 | 35.19 | 5.35E-07 | 2.22E-04 |
| Signaling by high-kinase activity BRAF mutants | 4 | 0.13 | 31.37 | 1.20E-05 | 2.71E-03 |
| Oncogenic MAPK signaling | 4 | 0.3 | 13.39 | 2.71E-04 | 2.42E-02 |
| MAP2K and MAPK activation | 4 | 0.14 | 28.16 | 1.77E-05 | 3.40E-03 |
| Signaling by RAF1 mutants | 4 | 0.15 | 27.45 | 1.95E-05 | 3.23E-03 |
| Post-translational protein phosphorylation | 10 | 0.39 | 25.66 | 1.39E-11 | 3.47E-08 |
| Signaling downstream of RAS mutants | 4 | 0.16 | 24.4 | 3.00E-05 | 4.67E-03 |
| Signaling by RAS mutants | 4 | 0.16 | 24.4 | 3.00E-05 | 4.15E-03 |
| Paradoxical activation of RAF signaling by kinase inactive BRAF | 4 | 0.16 | 24.4 | 3.00E-05 | 4.39E-03 |
| Signaling by moderate kinase activity BRAF mutants | 4 | 0.16 | 24.4 | 3.00E-05 | 3.93E-03 |
| Regulation of Insulin-like Growth Factor (IGF) transport and uptake by Insulin-like Growth Factor Binding Proteins (IGFBPs) | 10 | 0.45 | 22.14 | 5.43E-11 | 6.76E-08 |
| Platelet degranulation | 9 | 0.46 | 19.45 | 1.54E-09 | 1.28E-06 |
| Response to elevated platelet cytosolic Ca2+ | 9 | 0.48 | 18.72 | 2.12E-09 | 1.32E-06 |
| Signaling by BRAF and RAF1 fusions | 4 | 0.23 | 17.16 | 1.09E-04 | 1.14E-02 |
| Integrin cell surface interactions | 5 | 0.31 | 16.34 | 1.80E-05 | 3.21E-03 |
| Binding and Uptake of Ligands by Scavenger Receptors | 4 | 0.37 | 10.77 | 6.01E-04 | 4.83E-02 |

**Supplemental Table 3. $F_{ST}$ Values across ΔEP spectrum of values.**

Mean $F_{ST}$ rank value for UK10K SNPs in ΔEP bins for three population contrasts. Values are for SNPs that are in the UK10K sample and occur with at least 10 derived alleles in the pooled populations of the contrast. For each ΔEP SNP the observed $F_{ST}$ was ranked against that for control alleles of the same derived allele frequency.

| ΔEP | Africa vs Eurasia | Europe vs Asia | Great Britain vs Italy |
|---|---|---|---|
| -0.95 | 0.5615 | 0.5254 | 0.4913 |
| -0.85 | 0.5416 | 0.5249 | 0.5097 |
| -0.75 | 0.5388 | 0.5219 | 0.5112 |
| -0.65 | 0.521 | 0.5344 | 0.5106 |
| -0.55 | 0.5209 | 0.5211 | 0.5003 |
| -0.45 | 0.514 | 0.511 | 0.4985 |
| -0.35 | 0.4986 | 0.5182 | 0.5173 |
| -0.25 | 0.5178 | 0.4934 | 0.4936 |
| -0.15 | 0.5085 | 0.5328 | 0.4928 |
| -0.05 | 0.511 | 0.5223 | 0.4485 |
| 0.05 | 0.5076 | 0.5423 | 0.5211 |
| 0.15 | 0.5052 | 0.5544 | 0.4942 |
| 0.25 | 0.479 | 0.5825 | 0.5861 |
| 0.35 | 0.4844 | 0.532 | 0.5286 |
| 0.45 | 0.4902 | 0.547 | 0.4228 |
| 0.55 | 0.5889 | 0.5955 | 0.4614 |
| 0.65 | 0.5295 | 0.4961 | 0.4488 |
| 0.75 | 0.5159 | 0.6301 | 0.5388 |
| 0.85 | 0.4205 | 0.4204 | 0.4972 |
| 0.95 | 0.5693 | 0.4416 | 0.5675 |

**Supplementary Table 4. ΔEP measures for fixed and polymorphic alleles**

Based on maximum-likelihood rooting estimates of ancestral alleles (see Figure 1 for values based on Ensembl rooting). 95% confidence intervals on the mean, determined by bias-corrected bootstrap, are given in parentheses.

| | Negative ΔEP | | Positive ΔEP | |
|---|---|---|---|---|
| Measure | Fixed (%) | Polymorphic (%) | Fixed (%) | Polymorphic (%) |
| Count | 29348 (13.0%) | 195747 (87.0%) | 6941 (60.2%) | 4585 (39.8%) |
| Mean frequency | 1 | 0.025 | 1 | 0.096 |
| Mean Ancestral EP | 0.714 | 0.848 | 0.122 | 0.249 |
| Mean Derived EP | 0.052 | 0.017 | 0.602 | 0.515 |
| Mean ΔEP | -0.663 | -0.832 | 0.480 | 0.266 |

**Supplementary Table 5. Statistical power for detecting excess heterozygosity**

| Selection Coefficient | Probability of rejecting null hypothesis at false positive rate of 0.05 |
|---|---|
| 0.0 | 0.0485 |
| 0.0001 | 0.0545 |
| 0.0002 | 0.0560 |
| 0.0005 | 0.0615 |
| 0.001 | 0.0830 |
| 0.002 | 0.136 |
| 0.005 | 0.417 |
| 0.01 | 0.870 |
| 0.02 | 1.0 |
| 0.05 | 1.0 |
| 0.1 | 1.0 |

**Supplementary Figure 1. Distributions of derived polymorphism frequency in UK10K.**

Distribution of derived allele frequency for each ΔEP bin from –1 to +1 in 0.1 increments. Derived allele frequency ranges from singletons (1 copy of the derived allele) to 7241 copies (only one copy of the ancestral allele). The majority of sites are found at low frequencies across all bins.

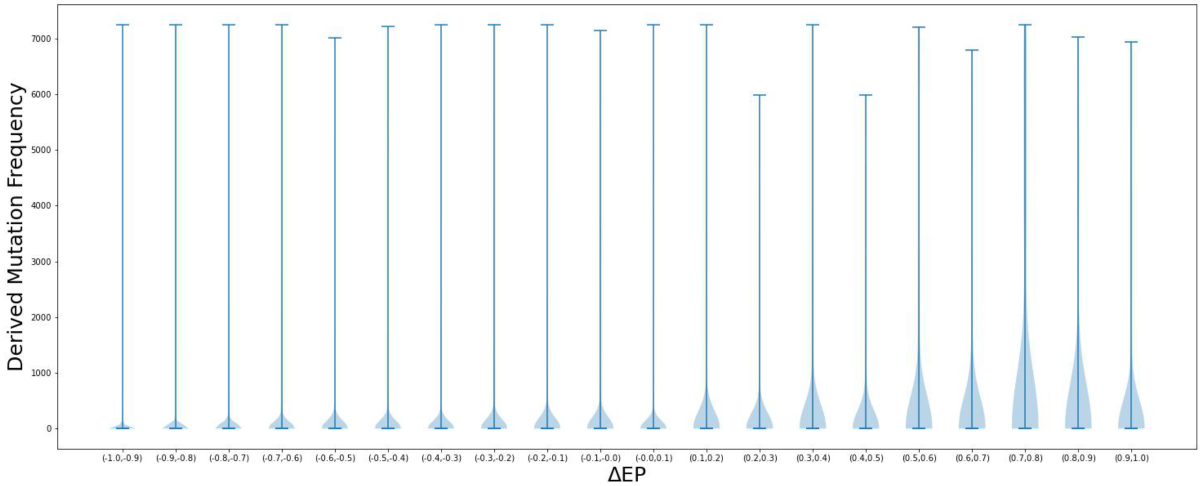
